# PDMQ-Protein Digestion Multi Query software tool to perform *in silico* digestion of protein/peptide sequences

**DOI:** 10.1101/014019

**Authors:** Reka Haraszi, Csongor Tasi, Angela Juhasz, Szabolcs Makai

**Affiliations:** independent consultant; Agricultural Institute, Centre for Agricultural Research, Hungarian Academy of Sciences (ARI CAR HAS), Brunszvik u. 2., Martonvásár 2462, Hungary

**Author notes:** corresponding author, /317. Currently at Campden BRI, Chipping Campden, UK.

## Abstract

**Motivation:** *In silico* enzymatic digestion tools mostly can be used for digestion of single sequence query, which means a significant limitation in their utility when a number of sequences need to be processed. The other limitation of these applications is the selection options of restriction enzymes that are usually allow only simultaneous digestion. Non-conventional proteins such as cereal prolamins require multi-enzyme multi step digestion, and for cereal proteomics experts this type of application is missing.

**Results:** **PDMQ, Protein Digestion Multi Query** application was developed having multi query and multi enzyme options and that way can be customized for any digestion protocol.

**Availability and implementation:** PDMQ is implemented in C# using the .NET framework and can be downloaded from http://www.agrar.mta.hu/_user/browser/File/bioinformatics/ProteinDigestion_v0_0_0_15.rar

## 1 INTRODUCTION

In proteomics studies, protein/peptide sequences are cut using restriction enzymes that cleave at specific sites of the amino acid chain. Today, with the spread of bioinformatic tools, a few *in silico* digestion freeware can be found on the internet like ExPaSy Peptide cutter (Gasteiger et al. 2005), mMass (http://www.mmass.org/), Protein Digestion Simulator (PDS) (http://omics.pnl.gov/software/ProteinDigestionSimulator.php) that provides a useful model for a “perfect” enzymatic digestion, which then is used for sequence identification in proteomics studies. Most of these tools can be used for single digestion, which means a significant limitation in their utility when a number of sequences need to be processed. The other limitation of these applications is the selection options of restriction enzymes. In protocols that are not using (only) trypsin or applying multiple enzymes; the performance of *in silico* digestion is quite problematic with the currently available software. Generally, application specific software is the ideal; therefore we developed and introduce a new application, the **PDMQ, Protein Digestion Multi Query** which contains multi query as well as multi enzyme options and that way can be customized for any digestion protocol (Figure 1).

**Figure 1.**
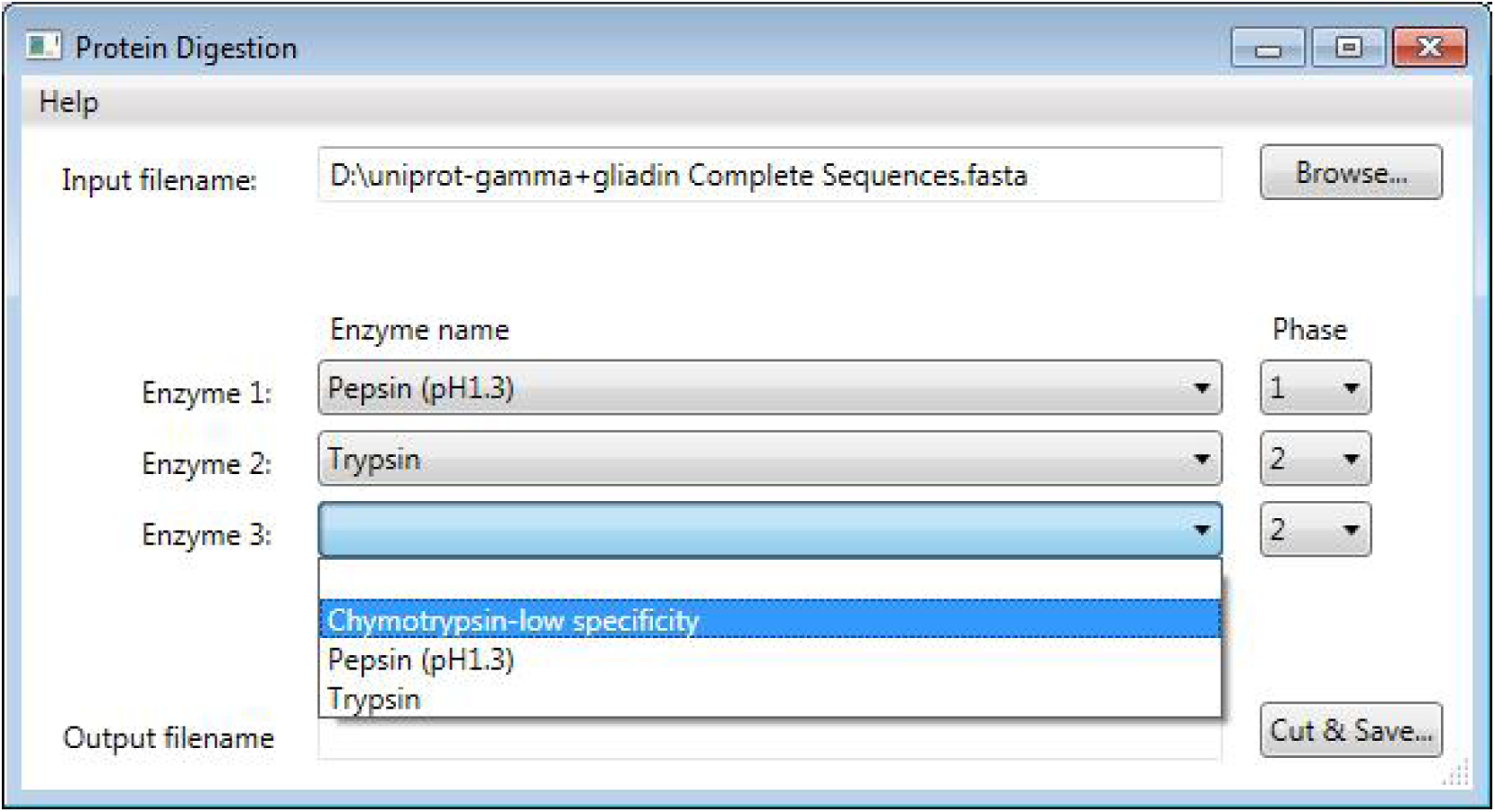
PDMQ main window shows the input file, the selection of three enzymes, pepsin for the first step, trypsin and chymotrypsin for simultaneous application for the second step.

## 2 RESULTS AND DISCUSSION

The unique feature of the software is the combination of the option for multi query input and that the enzymes can be applied in sequential order and/or simultaneously. For example, in a digestion protocol (Sealey-Voyksner et al. 2010) for gluten proteins, there is a two step protocol which applies pepsin in the first step and two other enzymes (trypsin and chymotrypsin) in the second step. PDMQ is able to perform the *in silico* analysis of a set of sequences with respect to the order of the applied enzymes (PEP 1, TR 2, CTR 2).

In the current available form of PDMQ, cleavage is possible of protein/peptide sequences of any length and with three enzymes: PEP (pepsin-pH 1.3), TR (trypsin) and CTR (chymotrypsin-low specificity). The digestion algorithm is identical to the ExPaSy Peptide cutter and was implemented in C#. During development, Peptide cutter was used to validate PDMQ. Input sequences are accepted in fasta, csv and txt formats. Results are given in tab separated values as a txt file keeping the format of the input file (e.g. fasta) and can be converted to a table containing the input sequence, sequence identifier, its length and mass, cleavage enzymes, cleavage positions, resulted peptide length and sequence(s) indicating their order in the digestion protocol, unrecognized amino acids and average mass [M^+^] of resulted peptides. The column of unrecognized amino acids notifies the user that these cases need to be treated manually and these amino acids are not considered in the mass parameter. Contrary, e.g. PDS [3] considers amino acid X with 113 Da in the average mass of a peptide but we have found it safer to let the user decide of the substitution of X and define the mass accordingly and manually.

### Cleavage algorithm of enzymes and their combinations

The enzymes can be applied in three different combinations:

1. single cleavage
2. simultaneous cleavage with different enzymes
3. subsequent cleavage with different enzymes

#### Single cleavage algorithm

PDMQ establishes cleavage sites by screening the input sequence from the N-terminal to the C-terminal (or in case of peptides in the order of the input) and investigates if each sites fulfill the cleavage criteria or not. Within an algorithm cycle, each and every cleavage is done as the total chain was cut, therefore it may happen that screening again the resulted sequence with the same enzyme, that sequence will be cleaved again at other sites according to other cleavage rules of the applied enzyme.

One or more cleavage rules and/or exception rules belong to an enzyme. An enzyme cleaves at a site when at least one cleavage rule and no exception rule apply. Cleavage criteria are considered to be valid, if all rules for the surrounding amino acids are valid, too. These rules concern the presence/absence of amino acids in relative positions to the cleavage site in both directions. A cleavage rule of an enzyme is the sum of elementary rules related to given positions. It includes eight elementary positions per one rule, four-four positions right before and after the cleavage site. Consequently, a cleavage rule only applies if all elementary rules apply in all the eight positions.

A set of amino acids belong to each elementary rule. An elementary rule can be either forward or inverse type. A forward elementary rule investigates the presence of an amino acid, from the defined set, in the given position and an inverse elementary rule investigates the absence of that.

In case of a forward rule, the elementary rule applies if an amino acid in the relative position related to the cleavage site (in the relative position of the elementary rule) is present in the set of amino acids belong to the elementary rule. In contrary, the absence of this amino acid gives validity for the inverse rule. If the set of amino acids is empty, then the elementary rule always applies.

If the elementary rule finds an amino acid X, it is never found in the defined amino acid set, therefore the forward elementary rule never applies but the inverse elementary rule always applies. If the relative position of the elementary rule is out of the input sequence (in case of the beginning and the end of the sequence), the forward rule is never applied and the inverse rule is always applicable.

#### Algorithm for simultaneous cleavage with different enzymes

The cleavage process is identical to the single cleavage algorithm with the only difference that each and every cleavage site is investigated according to rules of more enzymes. Resulted sequences contain all cleaved sequences obtained as a result of the simultaneous application of all used enzymes.

#### Algorithm for subsequent cleavage with different enzymes

The first step of this process is identical to the single cleavage algorithm, then using each resulted sequence as an input, is a subject of a second cleavage by the defined second enzyme according to the rules described in the single cleavage algorithm. In case of more enzymatic digestion steps this process is repeated always using the resulted sequences as input.

